# A dynamically-structured matrix population model based on renewal processes for accumulative development under variable environmental conditions

**DOI:** 10.1101/2021.02.04.429697

**Authors:** Kamil Erguler, Jacob Mendel

## Abstract

Arthropod vectors are responsible for the transmission of pathogens in humans and other species. The transmission rate depends on the size and activity of the vector population, factors which are in turn strongly affected by environmental conditions. Therefore, in order to develop realistic representations of vector population dynamics, it is necessary to properly account for the impact of a changing environment on the duration of life processes. Here, we use a pseudo-stage-structured population to model the accumulative process of development as a renewal process with a variable rate that depends on the environment. We incorporate this into sPop, formerly an age-structured population dynamics model. This framework allows the modeller to represent realistic life stage durations by choosing from three alternative probability schemes: an Erlang distribution, a Pascal distribution, or a fixed duration, while enabling the model to respond appropriately to variations in stage duration characteristics. Using this approach, we demonstrate that introducing random variation into the environmental conditions, which results in fluctuating development rates, on average decreases the time required for stage completion. An exception to this is an already optimum development rate being perturbed by noise towards a less efficient course. The proposed framework is suitable for performing inverse modelling with data collected from highly variable environmental conditions, the results of which can be used to develop realistic climate-driven population dynamics models.

## Introduction

Mathematical models of structured populations have been used extensively in population ecology (1–3) and infectious disease epidemiology (4). Recently, they have been used to predict habitat suitability and vector abundance in response to climate and environmental change (5–7).

Population heterogeneity as a result of the variation in the rate of a biological process, such as development or infectious period, may introduce a substantial time delay and play an important role in the emerging dynamics of a population (8). Numerous methods have been proposed to represent developmental heterogeneity. According to the way they structure a population, two main categories emerge: age-structured (1, 8–10) and pseudo-stage-structured models (4, 8, 11, 12).

In an age-structured model, individuals are grouped into a set of non-overlapping contiguous age-classes, each with a well-defined time interval. Members of the same class are considered identical, and life processes, such as survival and development, apply to each class in an age-dependent manner. Examples of age-structured models include the matrix population model of Leslie (1, 13), the integro-differential equations approach of Hethcote and Tudor (9, 10, 14, 15), the age-within-stage model of Caswell *et al*. (16), and the time-to-go formulation of Getz and Dougherty (17).

In certain age-structured models, the time spent in a particular life-stage (such as egg or larva in the case of modelling arthropods) is represented with a probability distribution (9, 10, 13–15, 17). These models invoke a survival function (or equivalently a hazard function) to determine the remaining time to stage completion. Here, we refer to this as the method of hazards. A major advantage of this method is its relative flexibility in representing various stage duration distributions.

In a pseudo-stage-structured model, individuals at various degrees of development are grouped into a set of sub-stages, which may or may not correspond to tangible life-stages. The former case, where the sub-stages are defined according to certain biological principles or discernible developmental features, is known as stage-structured modelling (2, 3). In comparison, the function of pseudo-stages is to accumulate an internal process, too complex to model or outright intangible, and thus control the time taken to develop all the way to the next life-stage (4, 8, 11). Here, we refer to this method as the method of accumulation. Insects are poikilotherms, which are strongly affected by variations in environment, especially in temperature (18–23). It is important for the predictive models of insect population dynamics to incorporate realistic development time distributions, which behave reliably under variable environmental conditions. Numerous modelling frameworks have been implemented specifically to tackle the problem of predicting the impact of environmental variation on insect populations, including the degree-day approach (24–26), matrix population models (with environmental perturbations (3, 27, 28) and temperature forcing (6, 29)), and ordinary (30, 31), delay (32, 33), and partial differential equations (34). However, only a subset of these incorporate realistic development time distributions (6, 29, 31).

Despite their flexibility in incorporating development time distributions, age-structured models are limited in their ability to model a scenario in which the survival function is time-varying as a result of fluctuations in environmental conditions. Recently, Erguler *et al*. have developed a dynamic age-structured model, which allows the hazard function to change with the environment (13, 29, 35). The change is imposed upon the existing age-structure under an assumption that the survival function lacks memory; that at any given time, the survival function for an individual depends on its age and on the current environmental conditions, but not on the historical conditions. However, the biological implications of this approach have not been tested thoroughly.

Pseudo-stage-structured models can accomodate variations in the development time distribution to a certain extent. A common technique for developing pseudo-stage-structured models is to employ a series of identical sub-stages with exponentially-distributed dwell times to yield an ordinary differential equations (ODE) system with an Erlang-distributed time for development from one life-stage to the next (also known as the linear chain trick) (8, 12). This approach has been generalised for a broader set of development time distributions (12) and adapted for stochastic modelling (31, 36, 37). When using the linear chain trick, one can transform variations in the rate of transition through the sub-stages into variations in the Erlang distribution characteristics (8, 31). However, due to the fixed number of sub-stages, change in the ratio of the mean to standard deviation is not permitted.

Here, we develop an algorithm to implement accumulative development with a dynamic number of pseudo-stages. The framework we employ is a discrete-time matrix population model with a dynamic state vector and projection matrix. The approach allows for Erlang-distributed, Pascal-distributed, and fixed stage durations, and permits variation in both the number of pseudo-stages and stage transition rates. We incorporate this method into the sPop software package (13), and demonstrate that the method of hazards and the method of accumulation respond differently to environmental variation, with the latter yielding more realistic development times under variable environmental conditions. Using the method of accumulation, we investigate the impact of a fluctuating environment on development, and discuss its repercussions on assessing the impacts of climate change on vector-borne diseases.

## Methods

To implement the method of accumulation, we consider a structured population where development is a renewal process (38) with a dynamic probability of event arrival. While each independent renewal event corresponds to a fraction of development, individuals with identical development fractions are grouped together.

In this population, the number of independent renewal events arriving at each iteration (time step) is the discrete random variable *i* with cumulative density function *F* (*x, θ*) = Pr(*i* ≤ *x*), where *θ* = 𝔼(*i*) ^*−* 1^ is the interarrival time. By choosing an appropriate expression for *F* (*x, θ*), it is possible to obtain Erlang- or Pascal-distributed development times or to simulate a deterministic fixed-duration development process.

The corresponding fraction of development achieved with each event is *κ*, given by *κ* = 1*/k*, the target number of events before development, so the rate of accumulation per time step is given by the product *κi*. Under the specific restrictions of invariable *k* and deterministic dynamics, this method is equivalent to a matrix population model, as we show in Supplementary Note A. To expand from the matrix population framework, we define the development indicator, *q*, such that initially *q* = 0 and at each time step *q* increments by *κi*, with development occuring when *q* ≥ 1. The development indicator acts as the pseudo-stage label such that all individuals with the same value of *q* are considered identical with respect to development - they are in the same pseudo-stage.

The advantage of *q* over *i* as the development indicator is that *q* provides a standard measure and allows for both *k* and *θ* to change. However, being a real-valued indicator between 0 and 1, *q* potentially, with a highly variable *k*, leads to an infinite number of pseudo-stages. In the stochastic setting, individuals are randomly assigned to a specific pseudo-stage, so the number of non-empty pseudo-stages never exceeds the number of individuals in a population. In contrast, a population can be divided indefinitely in the deterministic setting to represent the expected fraction in each pseudo-stage, and thus, the number of non-empty pseudo-stages can grow without bound. Under such circumstances, an approximation may be imposed by limiting the precision of *q*, which effectively limits the maximum number of pseudo-stages and groups individuals of almost identical progress together.

In order to facilitate dynamic handling of the pseudo-stages, allow for intrinsic stochasticity, and accomodate variable interarrival times, we developed Algorithm 1 (implemented in C and available in sPop v.2.0 under the GPL 3.0 license on the GitHub repository https://github.com/kerguler/sPop2). The algorithm simulates fixed or variable development times in either deterministic or stochastic settings. The user selects a distribution from fixed, Erlang, or Pascal for the development time distribution, and inputs the desired mean and standard deviation for each time step. The algorithm includes a scheme for selecting appropriate values of *k, θ* at each time step to specify *F* (*x, θ*) for suitably distributed development times.

The development time distribution referred to as fixed corresponds to the canonical degree-day approach (39), where stage completion and development occurs after a fixed duration without any variability, triggered by the accumulation of a precise number of degree-days. The rate of accumulation of degree-days is allowed to change depending on environmental drivers.

The fixed development scheme is a deterministic process with a target development time of *k* steps. This needs to be an integer value as it corresponds to the number of renewal events needed to complete the process. At each time step, precisely 1 renewal event accumulates (*θ* = 1). With each event, *κ* = 1*/k* of progress is added to the development indicator *q*, and the particular stage completes when *q* ≥ 1 as outlined above. If the conditions change during a simulation, and so does *k*, the rate of accumulation also changes. This leads to an intermediate development time between the initial and final *k*.

The Pascal development scheme, on the other hand, is an accumulative process designed to yield a Pascal-distributed development time, a special case of the negative binomial distribution with an integer-valued stopping parameter, *k*. This can be thought of as the distribution of the number of failures before *k* successes in a series of identical Bernoulli trials with probability of success *p*. According to this, a success corresponds to a renewal event, a failure corresponds to 1 time step, and the number of failures, *i*.*e*. time steps, needed for the *k*^*th*^ success, *i*.*e*. renewal event, is a Pascal-distributed random number, which corresponds to the development time. Since the number of successes before a failure, *i*, is geometrically distributed with E(*i*) = 1*/p*, we can identify the probability of success as *θ*. As expected, *q* increases by *i/k* at each time step.

It is important to note that, as an indicator of time, at least one failure event is required to be the final outcome in the series of Bernoulli trials. For instance, the probability of accumulating *k* successes in *τ* = 2 time steps is equal to the probability of accumulating *r* = 1 failure before *k* successes, where the second failure is a given. Therefore, *F* (*τ, k, θ*) = *F* (*r, k, θ*), where *F* (*τ, k, θ*) is the cumulative density function of a Pascal distribution with shape *k* and probability of success *p*, and *r* = *τ* − 1. In order to simulate this process, we write the probability of obtaining *x* successes before 1 failure for each time step as

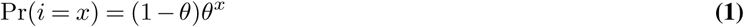

and the cumulative density function as

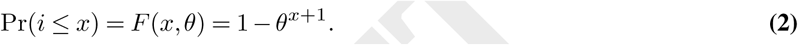

The final scheme we implemented is the Erlang, which yields Erlang-distributed development times, a special case of the gamma distribution where the shape parameter, *k*, is an integer. In this scheme, development time corresponds to the waiting time for the *k*^*th*^ renewal event in a Poisson process where interarrival times are exponentially distributed with rate *θ*. Consequently, the number of events arriving in a single time step, *i*, is a Poisson-distributed random number with rate 1*/θ*. The cumulative density function for the arrival rate can then be written as

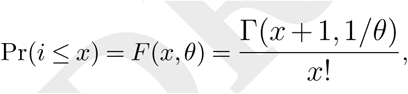

where Γ(*x, θ*) is the upper incomplete gamma function. As in the above cases, for each time step, *q* is incremented by *i/k*. The selection of a scheme is followed by iterating through the existing pseudo-stages, *i*.*e*. the existing groups of individuals with distinct *q* values. At first, all individuals have *q* = 0 so there is exactly one non-empty pseudo-stage representing a cohort. As time progresses, multiple values of *q* may appear representing other possible pseudo-stages. The algorithm also accommodates a structured initial population comprising a set of *q* values and associated sub-populations.

For the sub-population of individuals in pseudo-stage *q*_0_, the random variable *i* is iterated through all its possible values. For each value of *i, q* is incremented accordingly, *q*_*i*_ → *q*_0_ + *i/k*, and an appropriate fraction of the sub-population is allocated to *q*_*i*_ either deterministically, as the expected fraction of the population, or stochastically, by simulating a random fraction of the population, given the probability of *i*.

We exploited the hazard function, *i*.*e*. the probability of *x* events arriving conditional on the absence of fewer events,

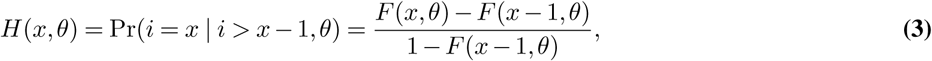

Where

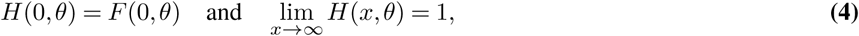

to accomplish the task of assigning a fraction of the population to each value of *i* in one pass.

Once the smallest value of *i* such that *q*_*i*_ ≥ 1 is reached, the remaining individuals, *m*, are flagged to complete development and the loop is terminated. As a result, for each *q* with an allocated population size of *n*, the algorithm generates one or more pseudo-stages, 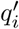, among which the population is distributed accordingly,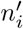.

It is important to note that the probability of *i* leading to *q >* 1 in a time step may become significant especially with a relatively large arrival rate. Overshooting of development, as we refer to such cases, will not prevent a population from completing development as required; however, may create bias should the process be followed by another stage of development. In such cases, the natural development process is interrupted by the aggregation of individuals with *q >* 1.

Overshooting can be minimised by extending the loop above *q* = 1 to maintain a record of all individuals with *q >* 1. In turn, the ones completing development form a structured population to be fed into the next stage of development. However, there may be cases where *q >* 2 in one step, which require multiple successive stages to utilise the accumulated development.

Consistent overshooting is a clear indication of a coarse time scale. However, due to the indirect time dependence of the renewal process, reducing the step size is not a trivial task. In the context of matrix population modelling, a time step is a unit based on which the survival and development processes are defined. Therefore, time scaling must involve a change in the desired duration and rate of these processes. For instance, daily survival probability, *λ*, can be scaled according to the relation,

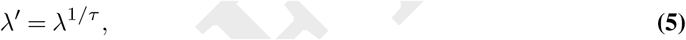

where the time interval is scaled down by a factor of *τ*. On the other hand, scaling the desired mean and standard deviation of an accumulative development process will render it appropriate for the new time scale,

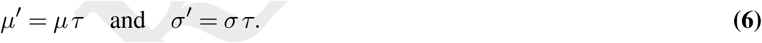

Time scaling results in a change in *k* with the fixed scheme, in *θ* with the Erlang, and in both with the Pascal scheme. We note that a finer time scale may result in a deviation from the target probability distribution when using the Pascal scheme due to the change in the shape parameter, *k*. Therefore, extra care must be taken when defining an accumulative process with the Pascal scheme.

### Algorithm 1

The accumulative development algorithm with discrete time steps. *I* is the indicator function, l*x*J indicates the largest integer value smaller than *x*, and l*x*l indicates rounding off to the nearest integer value.

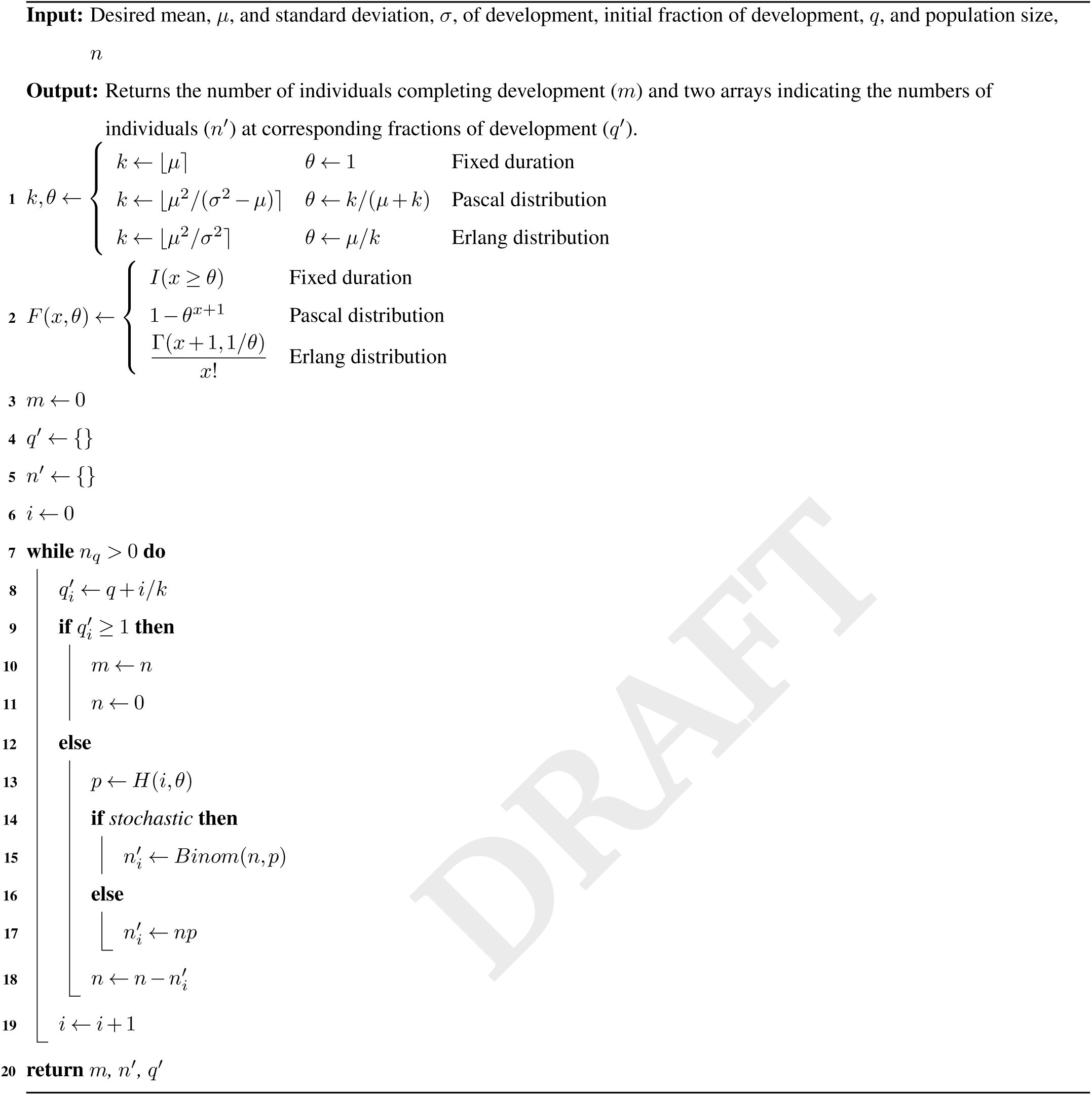

## Results and discussion

We developed Algorithm 1 to simulate an accumulative process as a stochastic renewal process in both deterministic and stochastic settings under three probability schemes: (i) fixed-duration, (ii) Pascal-distributed, and (iii) Erlang-distributed dwell time. The algorithm allows for variable dwell time characteristics, and thus, is aimed at representing realistic development times for natural populations under variable environmental conditions.

To demonstrate, we first simulated the temporal dynamics of development in a hypothetical population under constant conditions while excluding birth, death, and other biological processes. We created a small population of 10 individuals and imposed 20 ± 10 steps of development time, a mean of 20 steps with a standard deviation of 10. The emerging dynamics together with the impact of added stochasticity can be seen in Figure 1. As a result, we observed that the algorithm is capable of accurately simulating all three schemes as expected.

**Fig. 1.**
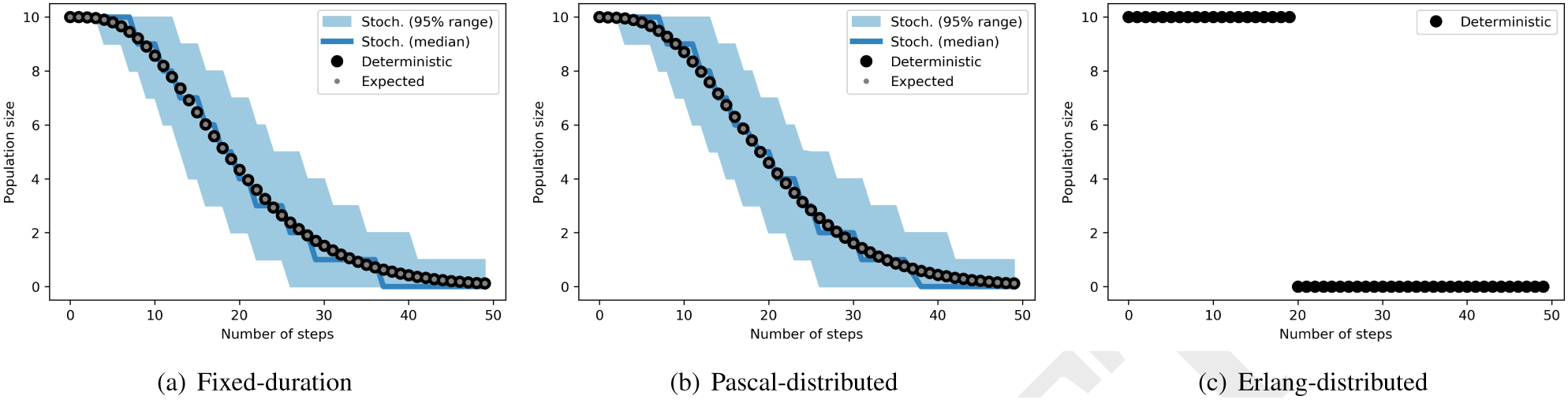
Three schemes for accumulative development. In (a), the fixed-duration scheme with 20 steps of development is shown. In (b), dwell time variability is incorporated in the form of a Pascal distribution with mean *µ* = 20 and standard deviation *σ* = 10 (corresponding to *θ* = 0.2 and *k* = 5). The same mean and standard deviation correspond to an Erlang distribution with *θ* = 5.0 and *k* = 4 in (c). The stochastic simulations were repeated 1000 times, and 95% output ranges are displayed as shaded areas.

We demonstrated the dynamics under variable conditions by introducing a drastic shift in development time — both in the shape and scale of the distribution — when only half the process was completed. In a deterministic scenario, we allowed a population of 100 individuals to develop for 20 time steps under conditions imposing 40 ± 5 steps of development, which corresponds to an Erlang distribution with *k*_0_ = 64 and *θ*_0_ = 0.625. Then, we switched the distribution to 20±5 steps, which requires *k*_1_= 16 and *θ*_1_ = 1.25.

We plotted the frequency diagram of the development indicator, *q*, for each time step in Figure 2(a). Note that the values of *q* are capped at just less than 1, since individuals with *q* ≥ 1 are removed from the population for development. Evidently, the switch affects the rate of change in *q*, but does not induce a shift in its position. Half the development was considered complete at the time of the switch as expected when employing the method of accumulation. Consequently, we observed a gradual stage transition with an overall development time of approximately 30±5 steps (Fig. 2(b), dark dots).

**Fig. 2.**
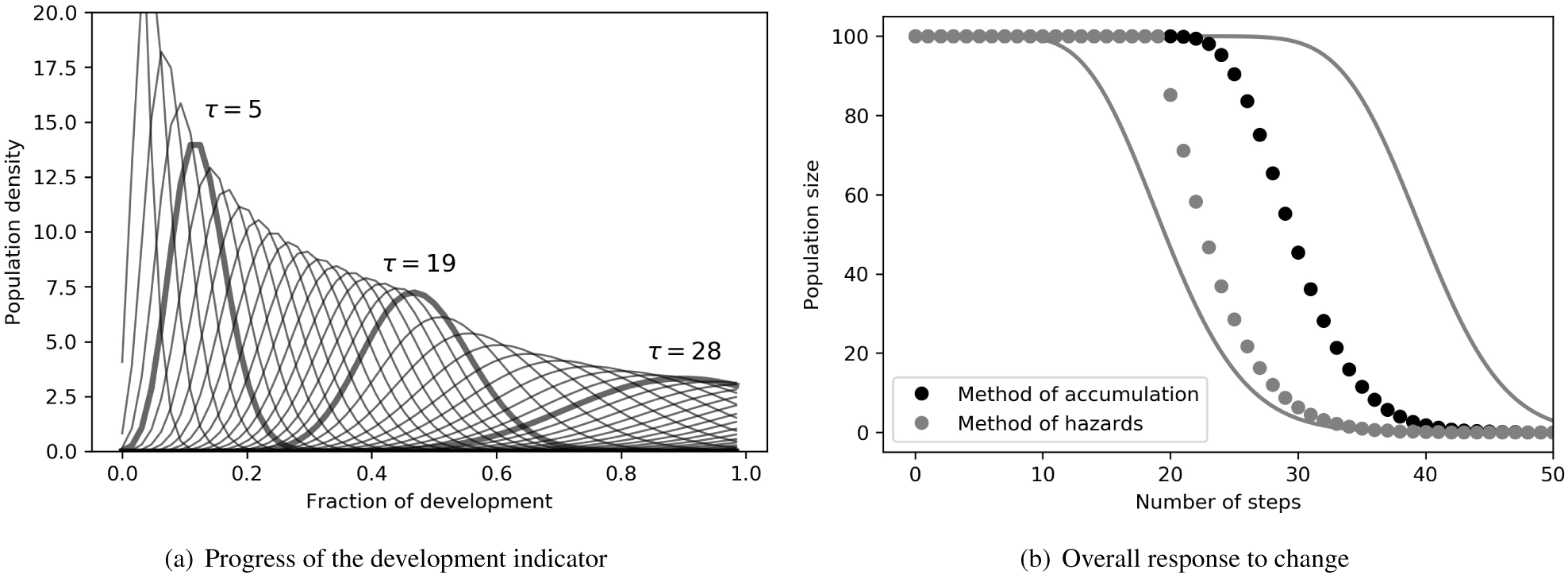
Response to change in development time. In (a), the frequency distribution of the development indicator, *q*, is shown for each time step. In (b), the number of developing individuals is shown as simulated by the accumulative development method (dark solid dots) and the age-dependent method of hazards (light solid dots). Solid gray lines mark the two target trajectories before and after the shift in development time.

In contrast, with the method of hazards, we observed a sharp transition at *τ* = 20 due to the strict age dependence (Fig. 2(b)). The switch at *τ* = 20 resulted in the majority of individuals reaching their target development age immediately. Evidently, the internal biological processes leading to development were overlooked, and the dynamics of stage transition was represented as being shaped by an external process. We argue that both representations might have appropriate applications for certain life processes; however, the method of accumulation is better suited for development.

Variation in the environment, especially in temperature, may affect the rate of development in arthropod vectors. Several, mostly non-linear, relationships have been proposed to represent the temperature dependency of insect development (22). A common feature in such proposals is the presence of an optimum temperature at which development is most efficient. An approximately linear trend is otherwise proposed for a wide range of temperatures provided that a clear distance is maintained from the optimum and the extreme conditions. To assess the impact of temperature variation in development, we considered the two alternative relationships illustrated in Figure 3(a); namely, the linear response to varying temperatures (shown in blue) and the optimum development efficiency being disrupted by variations in temperature (shown in red).

**Fig. 3.**
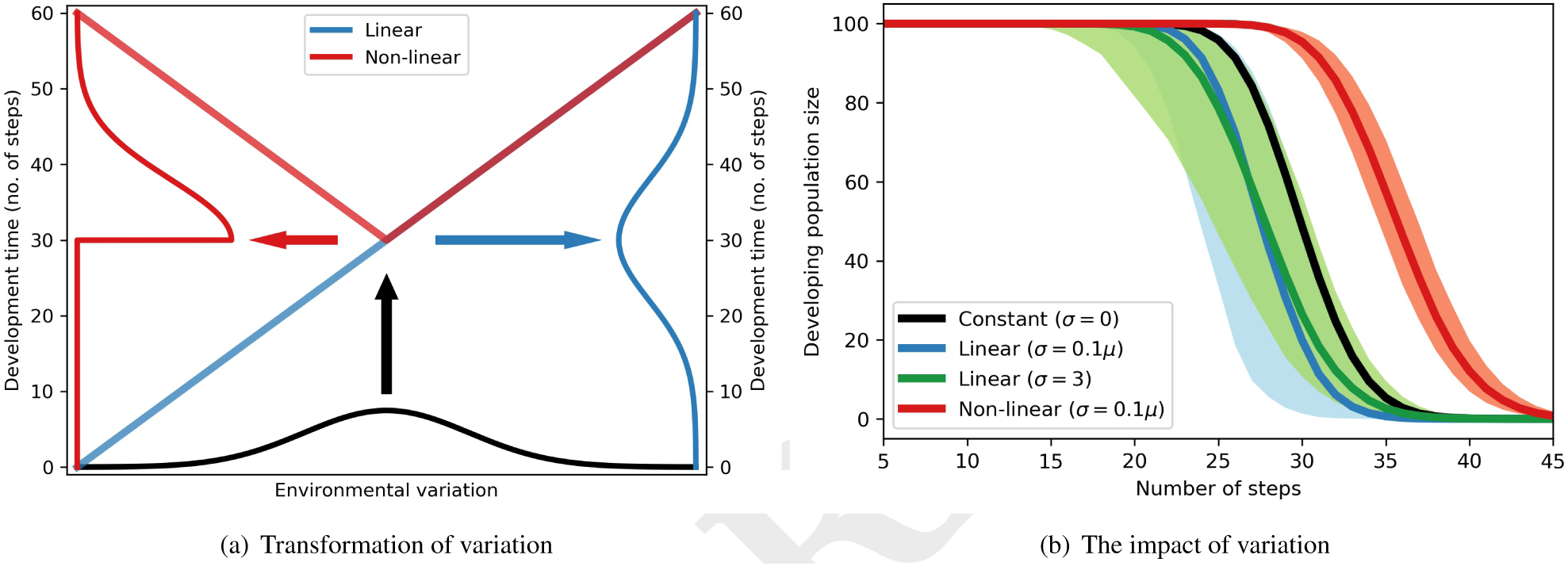
Accumulative development under environmental variation. In (a), relationship between environmental variation and average development time is shown. Solid lines indicate the frequency of noise and the average development time. In (b), the median (thick lines) and the 95% range (shaded areas) of development trajectories are shown for three variation scenarios. The thick black line indicates the case without environmental noise. The simulations were repeated 1000 times for each scenario.

We simulated a population of 100 individuals under three scenarios with reference to an Erlang-distributed development time of 30 ± 3 steps. In the first scenario, we assumed that the average development time, *µ*, varies linearly and independently for each time step in response to a Gaussian random number *ρ* ∼ ℕ(0, 8), and the standard deviation of the development time is proportional to the mean, *σ* = 0.1*µ*. We set the second scenario as identical to the first with the exception of a constant standard deviation, *σ* = 3 steps. In the final scenario, we set *ρ′* = |*ρ*| and *σ* = 0.1*µ* to test the impact of a non-linear response. We present indicative sets of development times for each scenario, in the form of corresponding Erlang distributions, in Figure S1, and the emerging development dynamics in Figure 3(b).

We note that in the second scenario, a different *k* is introduced with each step, thus the number of possible values of *q, i*.*e*. the number of pseudo-stages, increases rapidly in the deterministic setting. Due to the limitations in computational resources, we imposed an approximation by setting the precision of *q* to the nearest 1000^*th*^ decimal point, which effectively limits the maximum number of pseudo-stages to 1000 (see Methods). We demonstrated in Figure S2 that the approximation had a negligible impact on accuracy.

We compared the three scenarios against the reference case of no environmental variation (Fig. 3) and observed that the impact strongly depended of the relationship between temperature and development as expected. When development is already at an optimum efficiency, variation in temperature acts as a disturbance rendering the process less efficient and longer to complete (red shade in Fig. 3(b)). When it is away from the optimum, development time is reduced in response to noise due to the increase in the frequency of exposure to faster development rates (blue and green shades in Fig. 3(b)). This is in agreement with the experimental study of Wu *et al*. (2015), concerning *Megaselia scalaris*, which demonstrated that the temperature fluctuations between day and night boosted the development rate of the fly (25).

We observed that the two forms of standard deviation (proportional and fixed respectively in the first and second scenarios) gave rise to distinct characteristics of variation in the dynamics. With fixed *µ/σ*, the number of pseudo-stages is fixed, and so is the shape, *k*, of the development time probability distribution. Therefore, the variation observed in development time (blue and red shades in Fig. 3(b)) can be attributed to the variation in scale, *θ, i*.*e*. inverse of the rate of progress through each pseudo-stage. On the other hand, with fixed *σ*, the shape and scale change together keeping the ratio *k/θ* constant, which leads to a shift between a steep (when *k* is high) and a gradual response (when *k* is low). When *k* is consistently high, development takes longer to complete, but the process terminates with a steep curve. In contrast, consistently low *k* leads to shorter development times but a gradual termination curve, while very low *k* leads to transient spikes in the number of individuals completing development. Consequently, in accumulative development, low *k* early in the process has a large and lasting impact, which gives rise to the observed variability in Figure 3(b) (green shade).

Flexibility in defining the dwell time probability distribution provides an advantage for model and parameter inference, in which the duration and/or variability of the process cannot be determined *a priori*. The capacity to incorporate change in development rates during simulation also provides an advantage, and facilitates developing climate-driven population dynamics models for many vector species (6, 29).

While numerous modelling approaches concerning infectious disease dynamics require age-structured populations and associated delay in stage durations, such as the duration of the lag or infectious periods, a variable environment may not directly affect dwell times (4, 8). Even in cases where process characteristics are predetermined and flexibility is not a requirement, the extended sPop package introduced here offers an additional reliable high-performance tool for modelling structured population dynamics.

## Conclusions

Many species are exposed to large daily variations in temperature and rapid changes in microclimate as they grow and develop in their natural habitats. For many species, the rate of development is strongly associated with these drivers (40), and it is crucial to accurately represent the impact of these drivers in population dynamics models (41).

Here, we extended the age-structured population dynamics model sPop towards representing accumulative development with a dynamic pseudo-stage-structured population. The framework allows for the incorporation of realistic stage durations in population dynamics models while enabling to respond appropriately to variations in stage duration.

With the approach presented, the structure of a population is dynamically reconfigured into any number of sub-stages as needed to simulate two probability schemes — Erlang and Pascal distributions — and fixed duration. The number of sub-stages are limited only by the availability of computational resources. The algorithm allows for the distribution of development time (its mean, standard deviation, or type) to change during a simulation. As in the previous version of sPop, both deterministic and stochastic simulations are allowed.

Introduction of realistic stage durations in infectious disease models results in the increased complexity of transmission dynamics (8). We demonstrated that a variable stage duration further complicates the dynamics of a developing population by manifesting both the expected behaviour and the intrinsic stochastic variation. The effect must be accounted for in cases where stage duration is largely driven by environmental conditions, and especially when modelling climate-driven population dynamics of arthropod vectors.

The discrete-time matrix population framework possesses two main limitations, which may lead to numerical errors in simulation: (i) an integer value for *k* and (ii) discrete time steps. A restriction is imposed on certain combinations of mean and standard deviation, which lead to non-integer *k*. In such cases, the distribution is approximated by the nearest integer value. Future work will focus on relaxing this restriction and allowing for any non-integer value.

The numerical errors introduced by discrete time steps are largely subdued by the shortening of the steps, which can be informed by the expected duration of the process to be modelled. We argue that the loss in accuracy is minor in comparison to the ability to incorporate realistic dwell time distributions, flexibility in the number of sub-stages, and the performance gained.

Simulating realistic development times under varying environmental conditions is a vital requirement for developing predictive population dynamics models of disease vectors. The proposed approach is particularly appropriate for performing inverse modelling with data collected from highly variable environmental conditions.

## Supplementary Note A: The matrix population model

The matrix population modelling approach involves matrix algebra to project the state of a population to the next time point (3). According to this, a population is structured into a set of stages and/or within-stage age classes, and a projection matrix is constructed to describe the expected change in each stage/class during a time step.

Here, we derive the projection matrix for a special case of accumulative development with the deterministic assumption. By fixing the number of pseudo-stages to *k*, we eliminate the need to employ the development indicator *q*, and structure the population into *k* pseudo-stages, *n*_0_ … *n*_*k*−1_, and 1 additional stage, *u*, to receive the individuals completing development. By doing so, we map the development stages onto the projection matrix, and proceed to account for transitions from one pseudo-stage to multiple others.

To calculate the expected number of pseudo-stages one individual accumulates in one step, we use the cumulative density function describing the probability of accumulating *i* random renewal events in a single time step, *F* (*i*) = *F* (*i, θ*) (see Methods). For instance, an individual stays in a pseudo-stage with rate *F* (0), progresses to the next with rate *F* (1) − *F* (0), and skips one to reach the second pseudo-stage with rate *F* (2) − *F* (1). Consequently, the projection matrix, *M* (**n**, *τ*), can be written as

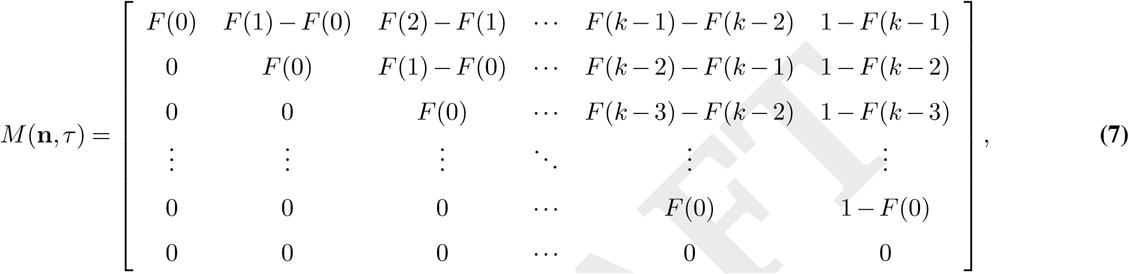

where **n** = [*n*_0_, *n*_1_, …, *n*_*k*−1_, *u*], *n*_*i*_ is the *i*^*th*^ pseudo-stage, and *u* is the subsequent development stage.

## Supplementary Note B: Environmental variation

**Fig. S1.**
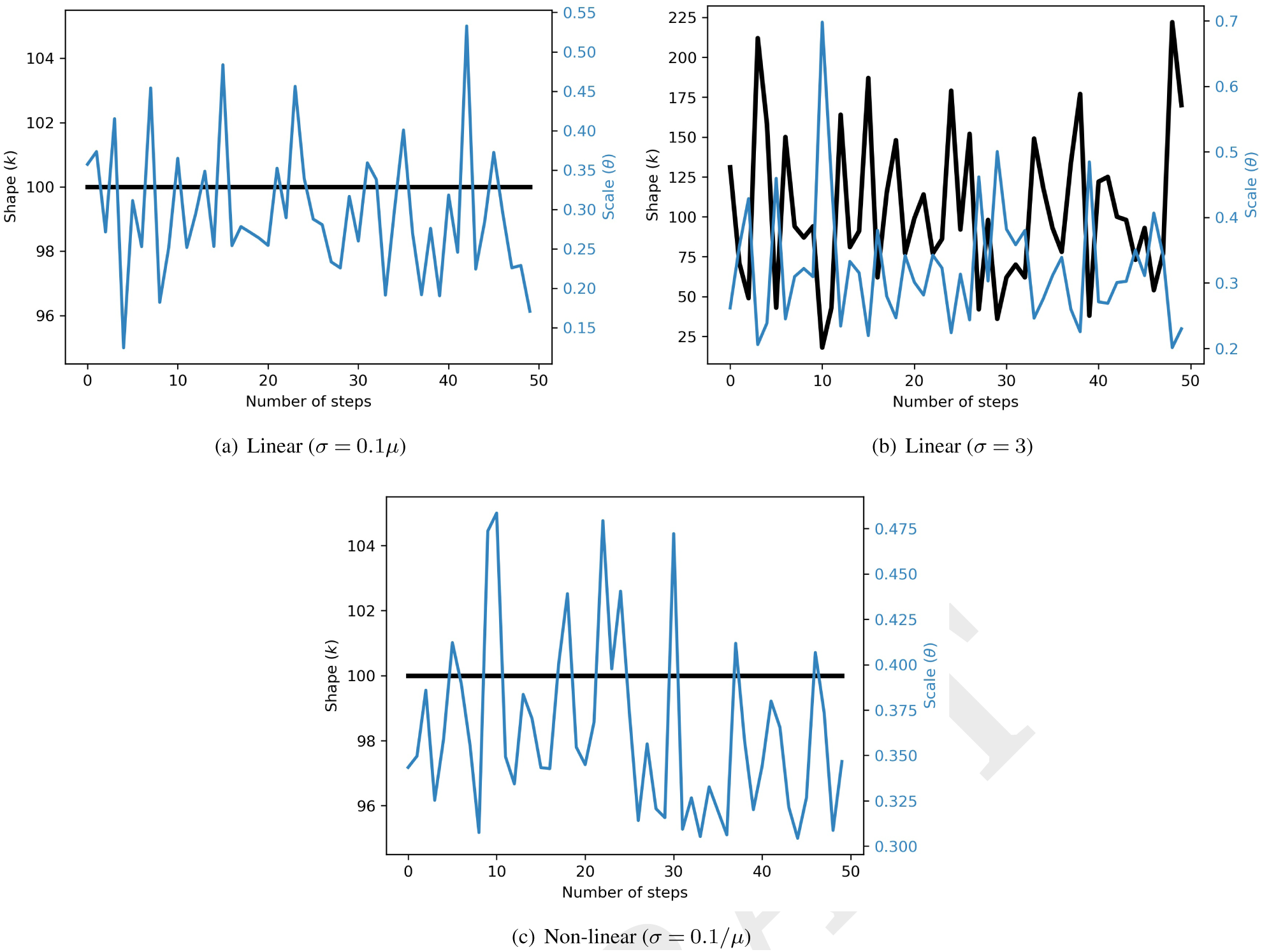
Environmental variation. A series of *k* (black) and *θ* (blue) are shown for the Erlang distributions corresponding to the development times sampled for three scenarios: (a) a linear change in *µ* with a proportional *σ*, (b) a linear change in *µ* with a constant *σ*, and (c) a non-linear change in *µ* with a proportional *σ*. See the main text for information on each scenario.

## Supplementary Note C: The approximation

**Fig. S2.**
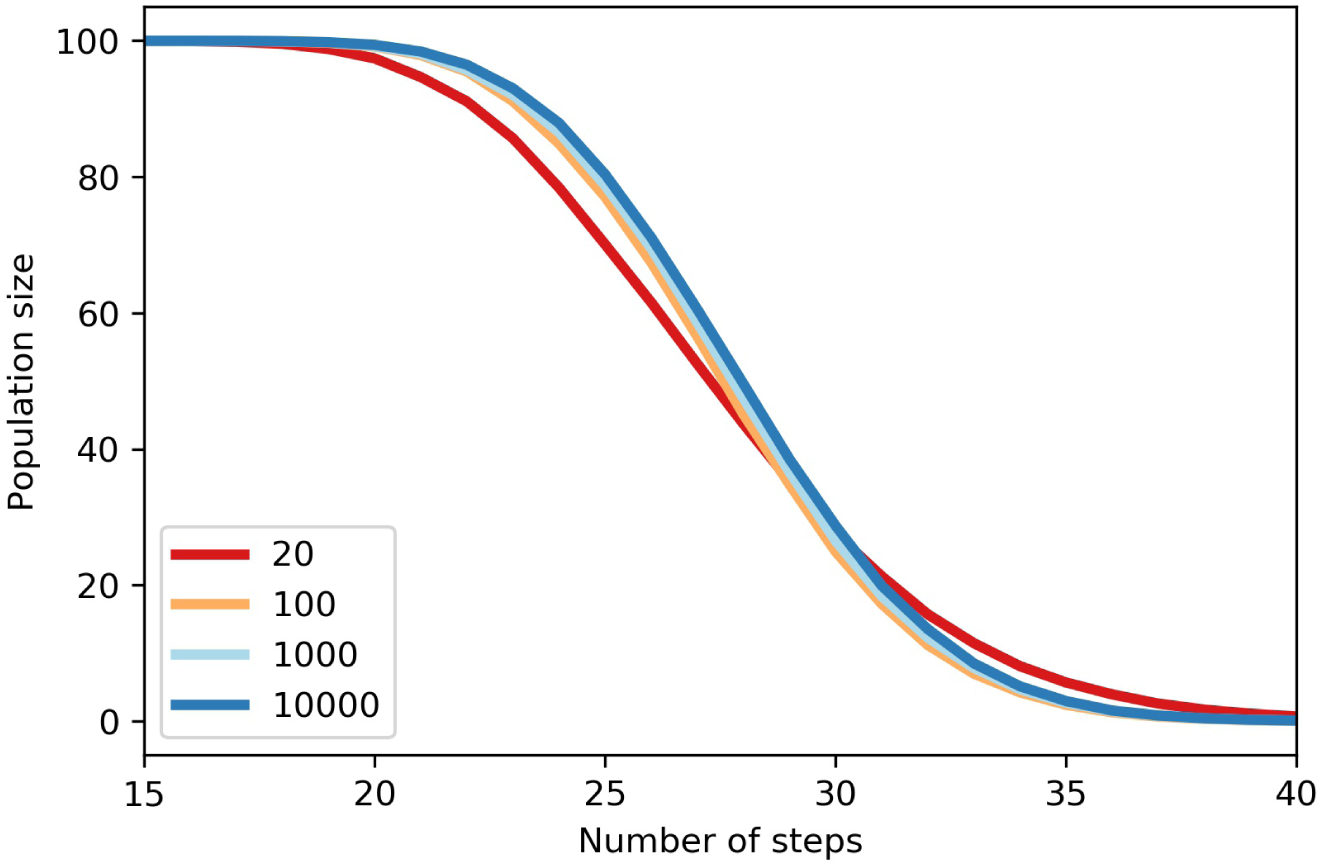
Limiting the number of pseudo-stages. The medians of development trajectories are shown for the variable temperature scenario (the second scenario with linear variation in *µ* and constant *σ*) when the number of pseudo-stages (plotted in different colours) are limited by tuning the precision of the development indicator *q* (see the main text).

## ACKNOWLEDGEMENTS

We thank Yiannis Proestos for valuable discussions. This publication has been produced within the framework of the EMME-CARE project which has received funding from the European Union’s Horizon 2020 Research and Innovation Programme, under Grant Agreement No. 856612 and the Cyprus Government. The authors declare that they have no competing interests. The sole responsibility of this publication lies with the authors. The funders had no role in study design, the decision to publish, or preparation of the manuscript. The funders are not responsible for any use that may be made of the information contained therein.

## Author contributions

KE conceived and supervised the study, developed the methods, performed the analyses, and wrote the manuscript. JM contributed to method development, data analyses, and to the final version of the manuscript.

## Notes

https://github.com/kerguler/sPop2

### Competing Interest Statement

The authors have declared no competing interest.

## Bibliography

1. P H Leslie. On the Use of Matrices in Certain Population Mathematics. Biometrika, 33(3):183–212, 1945. ISSN 00293970. doi:10.1093/nq/s3-XI.286.498-b.

2. L P Lefkovitch. The Study of Population Growth in Organisms Grouped by Stages. Biometrics, 21(1):1–18, 1965.

3. Hal Caswell. Matrix Population Models. Oxford University Press, 2001.ISBN 0878930965.

4. Roy M Anderson, Robert M May, and B Anderson. Infectious Diseases of Humans: Dynamics and Control. Oxford University Press, 1992. ISBN 019854040X.

5. Paul E Parham, Diane Pople, Céline Christiansen-Jucht, Steve Lindsay, Wes Hinsley, and Edwin Michael. Modeling the role of environmental variables on the population dynamics of the malaria vector Anopheles gambiae sensu stricto. Malaria Journal, 11:271, jan 2012. doi:10.1186/1475-2875-11-271.

6. Kamil Erguler, Stephanie E. Smith-Unna, Joanna Waldock, Yiannis Proestos, George K. Christophides, Jos Lelieveld, and Paul E. Parham. Large-Scale Modelling of the Environmentally-Driven Population Dynamics of Temperate Aedes albopictus (Skuse). PLOS ONE, 11(2):e0149282, feb 2016. ISSN 1932-6203. doi:10.1371/journal.pone.0149282.

7. Stéphanie Jenouvrier, Marine Desprez, Remi Fay, Christophe Barbraud, Henri Weimerskirch, Karine Delord, and Hal Caswell. Climate change and functional traits affect population dynamics of a long-lived seabird. Journal of Animal Ecology, 87(4):906–920, jul 2018. ISSN 00218790. doi:10.1111/1365-2656.12827.

8. Alun L. Lloyd. Destabilization of epidemic models with the inclusion of realistic distributions of infectious periods. Proceedings of the Royal Society of London. Series B: Biological Sciences, 268 (1470):985–993, may 2001. ISSN 0962-8452. doi:10.1098/rspb.2001.1599.

9. Herbert W. Hethcote and David W. Tudor. Integral equation models for endemic infectious diseases. Journal of Mathematical Biology, 9(1):37–47, mar 1980. ISSN 0303-6812. doi:10.1007/BF00276034.

10. M. J. Keeling and B. T. Grenfell. Disease extinction and community size: Modeling the persistence of measles. Science, 275(5296):65–67, 1997. ISSN 00368075. doi:10.1126/science.275.5296.65.

11. D.R. Cox and H.D. Miller. The Theory of Stochastic Processes. Routledge, sep 1977. ISBN 9780203719152. doi:10.1201/9780203719152.

12. Paul J. Hurtado and Adam S. Kirosingh. Generalizations of the ‘Linear Chain Trick’: incorporating more flexible dwell time distributions into mean field ODE models. Journal of Mathematical Biology, 79(5):1831–1883, oct 2019. ISSN 0303-6812. doi:10.1007/s00285-019-01412-w.

13. Kamil Erguler. sPop: Age-structured discrete-time population dynamics model in C, Python, and R [version 3; peer review: 2 approved]. F1000Research, 7:1220, 2018. ISSN 2046-1402. doi:10.12688/f1000research.15824.3.

14. Helen J Wearing, Pejman Rohani, and Matt J Keeling. Appropriate Models for the Management of Infectious Diseases. PLoS Medicine, 2(7):e174, jul 2005. ISSN 1549-1676. doi:10.1371/journal. pmed.0020174.

15. Scott Greenhalgh and Troy Day. Time-varying and state-dependent recovery rates in epidemiological models. Infectious Disease Modelling, 2(4):419–430, nov 2017. ISSN 24680427. doi:10.1016/j.idm.2017.09.002.

16. Hal Caswell, Charlotte de Vries, Nienke Hartemink, Gregory Roth, and Silke F. van Daalen. Age x stage-classified demographic analysis: a comprehensive approach. Ecological Monographs, 88 (4):560–584, nov 2018. ISSN 00129615. doi:10.1002/ecm.1306.

17. Wayne M. Getz and Eric R. Dougherty. Discrete stochastic analogs of Erlang epidemic models. Journal of Biological Dynamics, 12(1):16–38, jan 2018. ISSN 1751-3758. doi:10.1080/17513758.2017.1401677.

18. Ozge Erisoz Kasap and Bulent Alten. Laboratory estimation of degree-day developmental requirements of Phlebotomus papatasi (Diptera: Psychodidae). J Vector Ecol, 30(2):328–333,dec 2005.

19. H. Delatte, G. Gimonneau, A. Triboire, and D. Fontenille. Influence of temperature on immature development, survival, longevity, fecundity, and gonotrophic cycles of Aedes albopictus, vector of chikungunya and dengue in the Indian Ocean. Journal of Medical Entomology, 46(1):33–41, jan 2009. ISSN 0022-2585. doi:10.1603/033.046.0105.

20. Filiz Gunay, Bulent Alten, and Ergi Deniz Ozsoy. Estimating reaction norms for predictive population parameters, age specific mortality, and mean longevity in temperature-dependent cohorts of Culex quinquefasciatus Say (Diptera: Culicidae). Journal of Vector Ecology, 35(2):354–362, 2010. ISSN 10811710. doi:10.1111/j.1948-7134.2010.00094.x.

21. Oliver Brady, Michael Johansson, Carlos Guerra, Samir Bhatt, Nick Golding, David Pigott, Hélène Delatte, Marta Grech, Paul Leisnham, Rafael Maciel-de freitas, Linda Styer, David Smith, Thomas Scott, Peter Gething, and Simon Hay. Modelling adult Aedes aegypti and Aedes albopictus survival at different temperatures in laboratory and field settings. Parasit Vectors, 6(1):351, dec 2013. doi:info:pmid/24330720.

22. Pei-Jian Shi, Gadi V.P. P. Reddy, Lei Chen, and Feng Ge. Comparison of Thermal Performance Equations in Describing Temperature-Dependent Developmental Rates of Insects: (I) Empirical Models. Annals of the Entomological Society of America, 109(2):211–215, mar 2016. ISSN 0013-8746. doi:10.1093/aesa/sav121.

23. Giovanni Marini, Mattia Manica, Daniele Arnoldi, Enrico Inama, Roberto Rosà, and Annapaola Rizzoli. Influence of Temperature on the Life-Cycle Dynamics of Aedes albopictus Population Established at Temperate Latitudes: A Laboratory Experiment. Insects, 11(11):808, nov 2020. ISSN 2075-4450. doi:10.3390/insects11110808.

24. Brett S. Nietschke, Roger D. Magarey, Daniel M. Borchert, Dennis D. Calvin, and Edward Jones. A developmental database to support insect phenology models. Crop Protection, 26(9):1444–1448, sep 2007. ISSN 02612194. doi:10.1016/j.cropro.2006.12.006.

25. T.-H. Wu, S.-F. Shiao, and T. Okuyama. Development of insects under fluctuating temperature: a review and case study. Journal of Applied Entomology, 139(8):592–599, sep 2015. ISSN 09312048. doi:10.1111/jen.12196.

26. Mohammad Ali Mirhosseini, Yaghoub Fathipour, and Gadi V.P. Reddy. Arthropod development’s response to temperature: A review and new software for modeling. Annals of the Entomological Society of America, 110(6):507–520, 2017. ISSN 00138746. doi:10.1093/aesa/sax071.

27. V. Kaitala and E. Ranta. Is the impact of environmental noise visible in the dynamics of age-structured populations? Proceedings of the Royal Society B: Biological Sciences, 268(1478):1769–1774, 2001. ISSN 14712970. doi:10.1098/rspb.2001.1718.

28. Finlay Scott and Alastair Grant. Visibility of the impact of environmental noise: a response to Kaitala and Ranta. Proceedings of the Royal Society of London. Series B: Biological Sciences, 271 (1544):1119–1124, jun 2004. ISSN 0962-8452. doi:10.1098/rspb.2004.2710.

29. Kamil Erguler, Irene Pontiki, George Zittis, Yiannis Proestos, Vasiliki Christodoulou, Nikolaos Tsirigotakis, Maria Antoniou, Ozge Erisoz Kasap, Bulent Alten, and Jos Lelieveld. A climate-driven and field data-assimilated population dynamics model of sand flies. Scientific Reports, 9(1):2469, dec 2019. ISSN 2045-2322. doi:10.1038/s41598-019-38994-w.

30. Annelise Tran, Grégory L’ambert, Guillaume Lacour, Romain Beno\^\it, Marie Demarchi, Myriam Cros, Priscilla Cailly, Mélaine Aubry-Kientz, Thomas Balenghien, and Pauline Ezanno. A rainfall- and temperature-driven abundance model for Aedes albopictus populations. IJERPH, 10(5):1698–1719, may 2013. ISSN 16604601. doi:10.3390/ijerph10051698.

31. Sean L. Wu, Jared B. Bennett, Héctor M Sánchez C, Andrew J. Dolgert, Tomás M. León, John M. Marshall, Héctor M. Sánchez C., Andrew J. Dolgert, Tomás M. León, and John M. Marshall. MGDrivE 2: A simulation framework for gene drive systems incorporating seasonality and epidemiological dynamics. biorXiv, 2020. doi:10.1101/2020.10.16.343376.

32. W S C Gurney, R M Nisbet, and J H Lawton. The Systematic Formulation of Tractable Single-Species Population Models Incorporating Age Structure. Journal of Animal Ecology, 52(2):479–495, 1983.

33. R. M. Nisbet and W. S.C. Gurney. The systematic formulation of population models for insects with dynamically varying instar duration. Theoretical Population Biology, 23(1):114–135, 1983. ISSN 10960325. doi:10.1016/0040-5809(83)90008-4.

34. S Pasquali, L Mariani, M Calvitti, R Moretti, L Ponti, M Chiari, G Sperandio, and G Gilioli. Development and calibration of a model for the potential establishment and impact of Aedes albopictus in Europe. Acta Tropica, 202:105228, feb 2020. ISSN 0001706X. doi:10.1016/j.actatropica.2019.105228.

35. Kamil Erguler, N.L. Nastassya L. Chandra, Yiannis Proestos, Jos Lelieveld, G.K. George K. Christophides, and P.E. Paul E. Parham. A large-scale stochastic spatiotemporal model for Aedes albopictus-borne chikungunya epidemiology. Plos One, 12(3):e0174293, 2017. ISSN 1932-6203. doi:10.1371/journal.pone.0174293.

36. Alun L. Lloyd. Realistic Distributions of Infectious Periods in Epidemic Models: Changing Patterns of Persistence and Dynamics. Theoretical Population Biology, 60(1):59–71, aug 2001. ISSN 00405809. doi:10.1006/tpbi.2001.1525.

37. Fan Bai and Linda J.S. Allen. Probability of a major infection in a stochastic within-host model with multiple stages. Applied Mathematics Letters, 87:1–6, 2019. ISSN 18735452. doi:10.1016/j.aml.2018.07.022.

38. Kosto V. Mitov and Edward Omey. Renewal Processes. SpringerBriefs in Statistics. Springer International Publishing, Cham, 2014. ISBN 978-3-319-05854-2. doi:10.1007/978-3-319-05855-9.

39. R Bonhomme. Bases and limits to using “degree days” units. European Journal of Agronomy, 13:1–10, 2000.

40. Hans T. Ratte. Temperature and Insect Development. In Environmental Physiology and Biochemistry of Insects, pages 33–66. Springer-Verlag, 1984. doi:10.1007/978-3-642-70020-0_2.

41. Vadrevu Sree Hari Rao and Ravi Durvasula, editors. Dynamic Models of Infectious Diseases. Springer New York, New York, NY, 2013. ISBN 978-1-4614-3960-8. doi:10.1007/978-1-4614-3961-5.

